# Chromatin State Distinguishes Injury-Responsive from State-Stabilizing Transcriptional Programs in Hybrid Hepatocytes

**DOI:** 10.64898/2026.05.04.722673

**Authors:** Jacqueline A Brinkman, Fransky Hantelys, Jesse Raab, Adam D Gracz

## Abstract

Liver injury induces rapid transcriptional responses in hepatocytes, yet the chromatin features that distinguish injured hepatocytes from healthy hepatocytes remain poorly understood. Using an integrated functional genomics approach combining bulk RNA-seq, ATAC-seq, and CUT&Tag profiling of H3K27ac and H3K27me3, we define the transcriptional and chromatin landscape of *Sox9*-expressing hepatocytes, which exhibit gene expression consistent with both hepatocyte and biliary identity. Under homeostatic conditions, *Sox9*+ hybrid hepatocytes (HybHeps) are rare and confined to the periportal space, while chronic injury induces an expansion of *Sox9*+ metaplastic hepatocytes (MetHeps). We identify three classes of differentially expressed genes associated with injury-responsive, state-associated, or shared regulatory programs and demonstrate that these classes are governed by distinct chromatin mechanisms. Injury-responsive transcription is driven primarily by dynamic chromatin accessibility remodeling at NF-κB– and AP-1–enriched regulatory elements, while state-associated and shared programs are reinforced through selective H3K27ac and H3K27me3 modification with comparatively stable accessibility. Relative to conventional hepatocytes, HybHeps encode a permissive chromatin landscape at injury-responsive loci under homeostatic conditions, consistent with epigenetic priming that facilitates rapid inflammatory activation. Projection of mouse-derived gene programs onto a human liver single-cell atlas encompassing both healthy and diseased hepatocytes confirms that SOX9-expressing hepatocytes preferentially engage injury-associated inflammatory modules while attenuating hepatocyte metabolic identity programs. Together, these findings define a chromatin-based regulatory dichotomy between inflammatory responsiveness and hybrid hepatocyte cell state stability, providing mechanistic insight into how differentiated epithelial cells integrate inflammatory signals while preserving cell state.

## Introduction

The adult liver exhibits remarkable regenerative capacity, yet the molecular mechanisms that control hepatocyte plasticity during chronic injury are not well understood. In response to sustained inflammatory stress, hepatocytes undergo transcriptional remodeling that suppresses mature metabolic identity while activating stress-response and reparative programs^1,2^. Although such adaptive responses are essential for tissue repair, prolonged injury can promote stable alterations in cell state that contribute to fibrosis, cirrhosis, and hepatocellular carcinoma^3,4^. How inflammatory signaling interfaces with state-associated regulatory programs to shape durable hepatocyte state transitions remains an unresolved question.

Recent studies have identified a subset of hepatocytes marked by *Sox9* expression that exhibit features of both hepatocyte and biliary identity ^5^. These so-called hybrid hepatocytes (HybHeps) localize to the periportal niche and expand in number following injury^5–7^, suggesting a role in regeneration^5,7,8^. Upon chronic liver injury, HybHeps expand and acquire features of metaplastic biliary-like cells; we refer to this injury-induced *Sox9+* hepatocyte state as metaplastic hepatocytes (MetHeps) to distinguish them from the homeostatic HybHep population. However, it is not known whether HybHeps and MetHeps represent distinct cell states or a continuum of *Sox9*+ hepatocyte identity, and the chromatin regulatory logic that distinguishes injury-driven transcriptional activation from state-associated stabilization has not been systematically defined.

Here, we integrate RNA-seq and ATAC-seq profiling of *Sox9+* HybHeps and *Sox9+* MetHeps with RNA-seq, ATAC-seq, and CUT&Tag profiling of H3K27ac and H3K27me3 in *Sox9*− conventional hepatocytes (cHeps) and *Sox9*+ HybHeps. We define both the injury-responsive transcriptional program and baseline regulatory architecture of *Sox9*-associated hepatocyte subpopulations. We demonstrate that chronic injury induces a coordinated inflammatory transcriptional program driven predominantly by dynamic chromatin accessibility remodeling ^9^. In contrast, state-associated and shared transcriptional programs exhibit minimal accessibility changes but are selectively enriched for histone modification–based regulation. Notably, HybHeps encode a more open chromatin landscape at baseline relative to cHeps, with enrichment of H3K27ac at injury-responsive loci consistent with epigenetic priming for inflammatory activation^10^. Finally, we show that SOX9+ hepatocytes in human liver disease exhibit conserved inflammatory regulatory signatures, supporting the translational relevance of this regulatory framework^11^.

Our findings reveal that hepatocyte plasticity is maintained by two distinct but intersecting chromatin regulatory modes: rapid accessibility remodeling mediating acute inflammatory responses within *Sox9+* hepatocytes, and histone-based stabilization reinforcing state-associated transcriptional programs that distinguish HybHeps and cHeps. This dichotomy provides a mechanistic framework for understanding how transient injury responses are integrated into durable alterations in hepatocyte state.

## Results

### Injury induces an inflammatory transcriptional program in hybrid hepatocytes

To define the transcriptional response to chronic liver injury within the HybHep compartment, we performed RNA-seq on FACS-isolated Sox9^EGFP+^ hepatocytes from uninjured controls and mice subjected to 6 weeks of DDC (3,5-diethoxycarbonyl-1,4-dihydrocollidine) diet (n=4 per group), hereafter referred as injured mice (**Figure 1A**). Variance-stabilized *Sox9* counts were comparable between groups, validating consistent isolation of the HybHep population across conditions (**Supplemental Figure 1A**). Principal component analysis (PCA) of the 500 most variable genes revealed clear separation by injury status, with PC1 explaining 79% of the total variance (**Figure 1B**), indicating that DDC-exposure drives a dominant and reproducible transcriptional shift in *Sox9*+ hepatocytes. Differential expression analysis identified 1,190 genes significantly upregulated and 725 genes significantly downregulated following injury (FDR < 0.05) (**Figure 1C**), demonstrating extensive injury-associated remodeling of the MetHep transcriptome.

**Figure 1.**
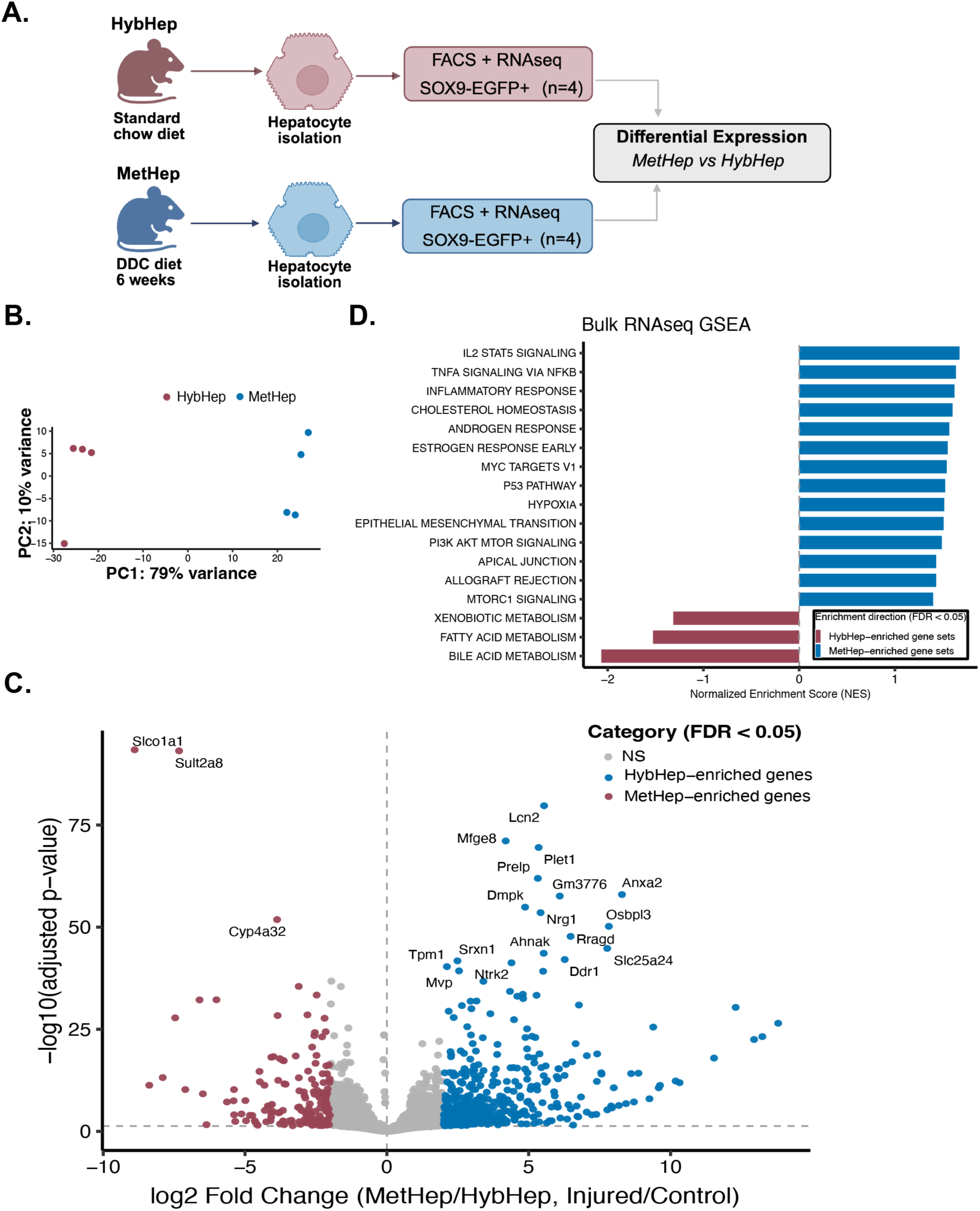
DDC diet induces a robust and reproducible transcriptional response in hybrid hepatocytes. **(A)** Experimental model. **(B)** Principal component analysis (PCA) of batch-corrected variance-stabilized (VST) RNA-seq counts. PC1 accounts for 79% of the total variance and separates samples by treatment condition, reflecting the injury response (n = 4). **(C)** Volcano plot showing log2 fold change and adjusted *P* values from DESeq2 differential expression analysis. A total of 1,190 genes were significantly upregulated and 725 genes were significantly downregulated in response to injury (Wald test with Benjamini–Hochberg multiple testing correction, adjusted *P* < 0.05). ApeGLM-shrunken log2 fold changes are shown. **(D)** Gene set enrichment analysis (GSEA) bar plot showing the most significantly activated and repressed gene sets following injury.

Gene set enrichment analysis (GSEA) revealed that injury-upregulated genes were strongly enriched for inflammatory and stress-response pathways, including TNFα signaling via NF-κB, IL-2/STAT5 signaling, and inflammatory response programs (**Figure 1D**). In contrast, injury-repressed gene sets were enriched for core hepatocyte metabolic programs including bile acid metabolism, fatty acid metabolism, and xenobiotic metabolism, indicating suppression of mature metabolic identity. Leading-edge analysis of the top injury-activated hallmarks demonstrated substantial overlap among the TNFα/NF-κB, inflammatory response, and IL-6/STAT3 gene sets (**Supplemental Figure 1B, left**), suggesting coordinated activation of a shared inflammatory regulatory module. On the other hand, leading-edge analysis of repressed metabolic hallmarks demonstrated extensive co-suppression of fatty acid, bile acid, and xenobiotic metabolism gene sets (**Supplemental Figure 1B, right**), consistent with broad attenuation of hepatocyte metabolic programs. Notably, signaling pathways commonly implicated in hepatocyte lineage reprogramming including, Notch, TGF-β, and Hippo, were not significantly enriched in MetHeps (**Supplemental Figure 1C**), suggesting that the transcriptional shift reflects inflammatory activation rather than reprogramming toward biliary epithelial identity. Together, these results define an injury-induced transcriptional state in MetHeps that is not present in HybHeps and is characterized by coordinated inflammatory pathway activation and concurrent attenuation of hepatocyte metabolic function.

### Injury-associated chromatin remodeling in MetHeps favors inflammatory transcription factor engagement

To determine whether transcriptional changes observed in MetHeps were accompanied by changes in chromatin accessibility, we performed bulk ATAC-seq on FACS-isolated *Sox9*+ hepatocytes from control and injured mice (n=4 per group). PCA of the 500 most variable regions revealed that DDC-treatment drives the majority of sample variance, consistent with RNA-seq results (**Supplemental Figure 2A**). Differential accessibility analysis identified 13,846 injury-gained and 13,273 injury-lost differentially accessible regions (DARs) (**Figure 2A**), indicating widespread chromatin remodeling in response to injury.

**Figure 2.**
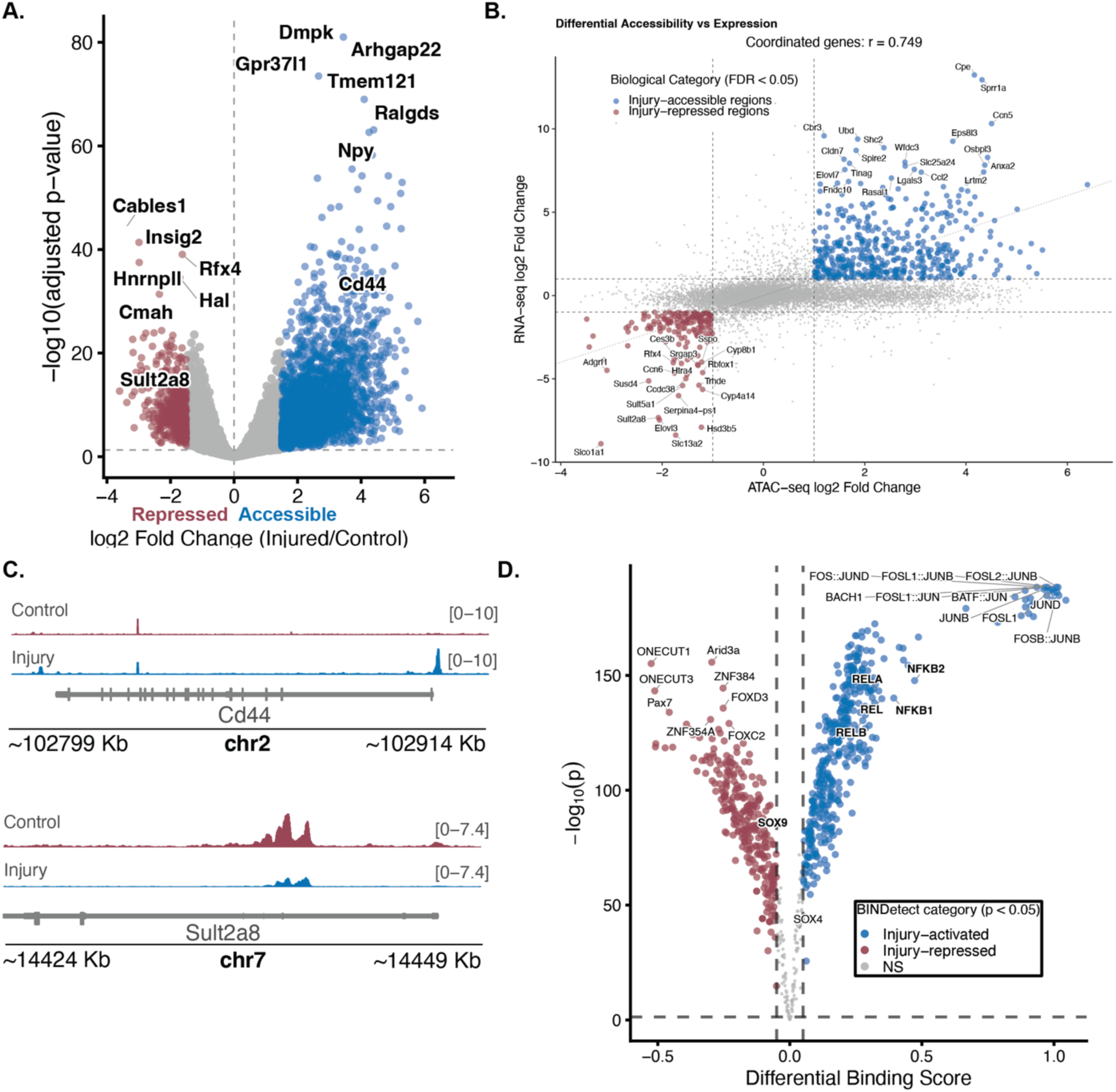
Bulk ATAC-seq reveals widespread and reproducible chromatin accessibility remodeling in response to injury. **(A)** Differential accessibility analysis identifies a near-equal distribution of injury-induced gains and losses in chromatin accessibility, with 13,846 regions significantly increased and 13,273 regions significantly decreased relative to control hybrid hepatocytes (Wald test with Benjamini–Hochberg correction, adjusted *P* < 0.1; n = 4). Gene labels indicate peak annotation to the nearest gene using ChIPseeker. **(B)** Pearson correlation analysis (r = 0.749) between differentially accessible regions and differentially expressed genes. For each gene, the region with the maximal ATAC-seq signal was used. **(C)** Representative genome browser tracks illustrating injury-induced accessibility changes at selected loci. ATAC-seq signal and representative tracks for *Cd44* and *Sult2a8*. Tracks represent mean signal across biological replicates (n = 4). **(D)** TOBIAS results of differentially accessible injury-responsive regions show AP-1 and NF-κB are predicted to gain binding, whereas the ONECUT transcription factors are predicted to lose binding.

Integration of ATAC-seq and RNA-seq datasets revealed a strong positive correlation between chromatin accessibility and gene expression changes (r=0.749, **Figure 2B**). Coordinated injury-activated genes clustered in the upper-right quadrant of the accessibility-expression space, whereas injury-repressed genes localized to the lower-left quadrant, demonstrating concordant chromatin and transcriptional remodeling. Genome browser visualization of representative loci illustrated these coordinated dynamics: *Cd44,* which is associated with biliary epithelial cells and MetHeps, gained chromatin accessibility and increased expression following DDC treatment, while the bile acid sulfotransferase *Sult2a8* showed reduced accessibility and decreased expression (**Figure 2C, Supplemental Figure 2B**). A subset of genes exhibited discordant or single-modality changes, suggesting additional layers of post-transcriptional regulation or distal enhancer-promoter interactions not fully resolved by bulk ATAC-seq (**Figure 2B)**.

To identify candidate regulatory drivers of injury-induced chromatin remodeling, we performed transcription factor footprinting analysis on injury-gained DARs using TOBIAS. Injury-gained DARs were strongly enriched for predicted inflammatory transcription factor DNA binding, including NF-κB family members (RELA, RELB, NFKB1, NFKB2), AP-1 family members (FOS, JUN, and related heterodimeric complexes), and STAT factors (**Figure 2D**). In contrast, motifs for hepatocyte lineage regulators including ONECUT and FOXD family members were enriched in regions losing accessibility, consistent with reduced occupancy at hepatocyte metabolic gene regulatory elements in the injured state. The predicted displacement of ONECUT1 and FOXD factors aligns with the suppression of bile acid, fatty acid, and xenobiotic metabolism gene sets observed by RNA-seq (Figure 1C), supporting that MetHeps exhibit both inflammatory activation and loss of lineage-defining transcription factor engagement. Although SOX9 and SOX4 motifs are present among injury-gained DARs and show statistically detectable differences in predicted occupancy between conditions (SOX9 Δscore = −0.11; SOX4 Δscore = +0.03), these effect sizes are small relative to the large positive binding score changes observed for AP-1 family members (FOS::JUNB Δscore = +1.05; FOSL2::JUND Δscore = +1.02) and NF-κB family members (NFKB2 Δscore = +0.47; RELA Δscore = +0.34, **Figure 2D**).

To determine whether inferred changes in transcription factor occupancy were reflected at the transcriptional level, we examined VST-normalized expression of representative inflammatory and state-associated regulators. NF-κB family members (*Rela, Relb*), AP-1 components (*Fos, Jun*), and *Stat3* showed maintained or increased expression in injured HybHeps, while the state-associated regulator *Onecut1* was significantly reduced (**Supplemental Figure 2C**). Broader profiling of AP-1, NF-κB, ONECUT, and STAT family members confirmed that inflammatory TF families were upregulated whereas *Onecut1* expression was selectively attenuated, consistent with a shift in the active regulatory landscape away from hepatocyte identity programs. These data suggest that chronic liver injury induces coordinated chromatin reorganization in MetHeps that favors enhanced accessibility at inflammatory regulatory loci and reduced accessibility at metabolic-associated loci, reinforcing the emergence of an inflammatory transcriptional state while attenuating hepatocyte identity programs.

### Hybrid hepatocytes encode a permissive enhancer landscape relative to conventional hepatocytes

To define the baseline regulatory architecture of HybHeps relative to cHeps, we performed bulk RNA-seq, ATAC-seq, and CUT&Tag profiling of H3K27ac and H3K27me3 under homeostatic conditions (**Figure 3A, Supplemental Figure 3A-C**). Transcriptomic comparison revealed subtle gene expression differences between HybHeps and cHeps (**Figure 3B**), confirming that these populations represent distinct functional states. HybHeps preferentially expressed genes associated with inflammatory responsiveness and biliary epithelial identity, including *Sox9*, *Spp1*, and *Arhgap10*, whereas cHeps exhibited stronger expression of canonical hepatocyte metabolic programs including cytochrome P450 family members and bile acid metabolism genes. These transcriptional differences were accompanied by a slight divergence in chromatin accessibility. Genome-wide ATAC-seq analysis identified 116 regions with increased accessibility and 17 regions with decreased accessibility in HybHeps relative to cHeps (**Figure 3C**). However, PCA for ATAC-seq and CUT&Tag profiling confirmed that HybHep identity drives a majority of differences observed for each assay, indicating that *Sox9*+ hepatocytes maintain a distinct chromatin landscape even in the absence of injury **(Supplemental Figure 3A-C**).

**Figure 3.**
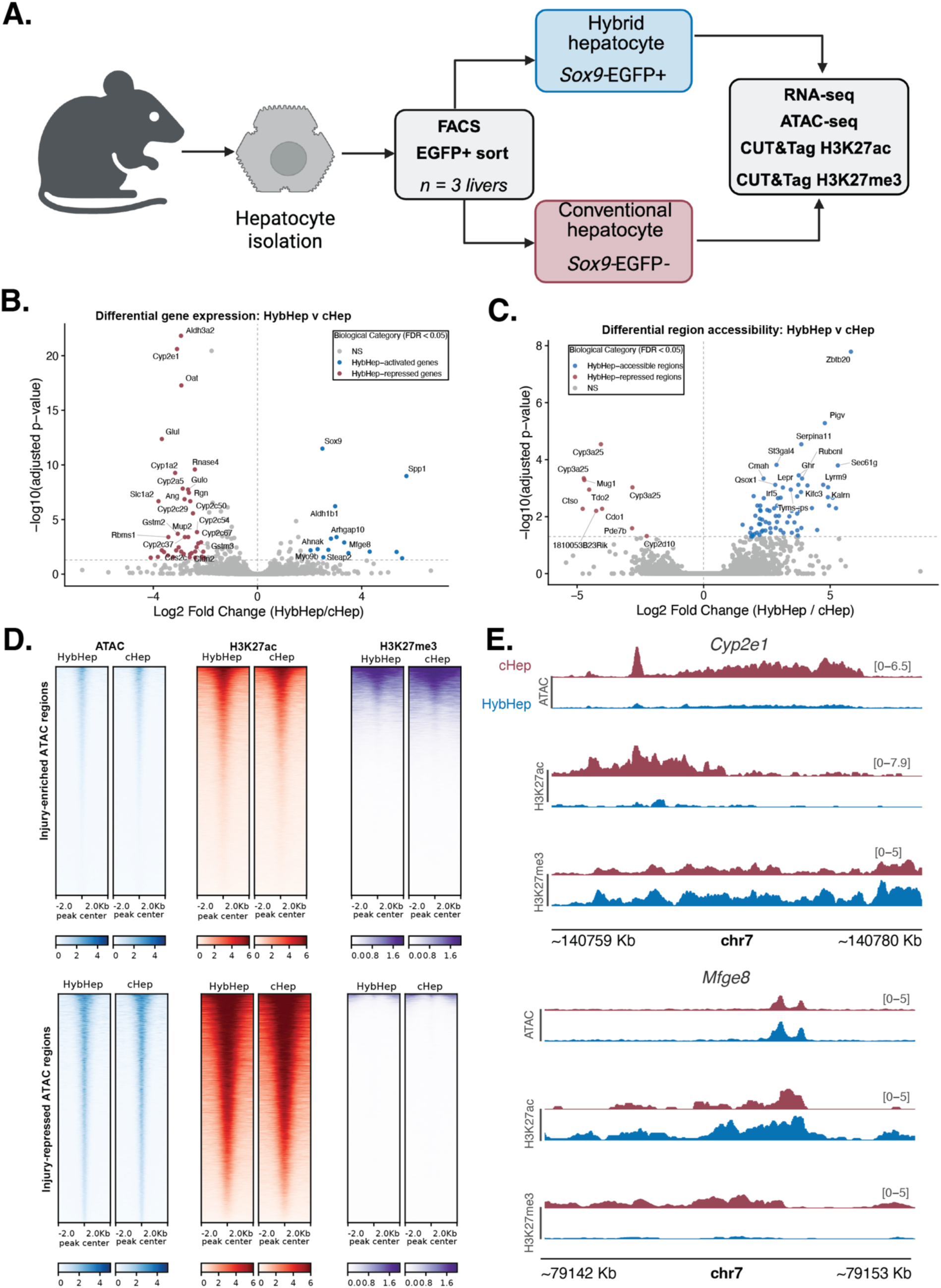
Hybrid hepatocytes encode a permissive enhancer landscape relative to conventional hepatocytes. **(A)** Experimental approach. **(B)** Volcano plot showing log2 fold change and adjusted p-values from DESeq2 differential expression analysis comparing HybHeps to cHeps. HybHep-activated and HybHep-repressed genes are highlighted (Wald test with Benjamini–Hochberg correction, adjusted P < 0.05). **(C)** Volcano plot of differential chromatin accessibility between HybHeps and cHeps. HybHep-accessible and HybHep-repressed regions are indicated (adjusted P < 0.1). **(D)** Tornado plots showing ATAC-seq, H3K27ac, and H3K27me3 signal centered ±2 kb from the midpoint of injury-gained (top) and injury-lost (bottom) accessible regions, shown for HybHep and cHep samples. Injury-gained regions are enriched for H3K27ac and depleted for H3K27me3 in HybHeps relative to cHeps, consistent with a permissive baseline enhancer configuration at injury-responsive loci. Injury-lost regions show the reciprocal pattern. Rows are sorted by ATAC signal in HybHeps. **(E)** Genome browser tracks at representative loci illustrating the multi-layered epigenetic differences between HybHeps and cHeps. *Cyp2e1* (left), a canonical hepatocyte metabolic gene repressed in both the state and injury comparisons, exhibits reduced chromatin accessibility, decreased H3K27ac, and elevated H3K27me3 in HybHeps relative to cHeps, consistent with a repressive chromatin configuration at hepatocyte identity loci. *Mfge8* (right), a HybHep-enriched gene, shows the reciprocal pattern of gained accessibility, H3K27ac enrichment, and reduced H3K27me3, consistent with an active enhancer configuration at this locus in HybHeps. Tracks display mean-normalized signal across biological replicates (n=3). Signal ranges are indicated in brackets.

To determine whether HybHep-specific chromatin features overlap with injury-responsive regulatory elements observed in MetHeps, we examined ATAC-seq, H3K27ac, and H3K27me3 signal at injury-gained and injury-lost DARs in HybHeps and cHeps. Injury-gained DARs showed enriched H3K27ac signal in HybHeps relative to cHeps and comparatively lower H3K27me3, consistent with a permissive enhancer configuration at these loci (**Figure 3D, top**). In contrast, injury-lost DARs displayed reduced H3K27ac and elevated H3K27me3 in HybHeps relative to cHeps, consistent with a repressive chromatin configuration at hepatocyte metabolic gene loci (**Figure 3D, bottom**). These cell-type differences in histone modification at injury-responsive regions were not observed to the same degree in cHeps, suggesting that the epigenetic pre-configuration of injury-responsive regulatory elements is a HybHep-specific feature (**Supplemental Figure 3D)**. Genome browser visualization at representative loci further illustrated these multi-layered epigenetic relationships: *Cyp2e1*, a canonical hepatocyte metabolic gene repressed in both HybHeps relative to cHeps and in injured relative to uninjured HybHeps, exhibited reduced accessibility, decreased H3K27ac, and increased H3K27me3 in HybHeps. *Mfge8*, a HybHep-enriched gene, showed the opposite pattern of gained accessibility, H3K27ac enrichment, and reduced H3K27me3 (**Figure 3E**). The correspondence between baseline H3K27ac enrichment and injury-gained DARs supports a model in which HybHeps encode a pre-primed enhancer landscape that facilitates rapid inflammatory activation and transition to a MetHep “state” upon injury.

To characterize the signaling programs associated with HybHep-enriched accessible chromatin, we performed GSEA on genes linked to HybHep-accessible ATAC-seq peaks. TNFα signaling via NF-κB was significantly enriched among genes associated with HybHep-DARs (NES = 1.34, padj = 0.005), while IL6/JAK/STAT3 signaling showed a positive but non-significant trend (NES = 1.30, padj = 0.065), and the broader hallmark inflammatory response gene set was not enriched (NES = 1.03, padj = 0.44). Critically, none of these gene sets were significantly enriched in the HybHep transcriptome under homeostatic conditions (TNFα/NF-κB NES = 1.05, padj = 0.92; IL6/JAK/STAT3 NES = −0.82; Inflammatory response NES = −0.85, **Supplemental Figure 3E**). The selective enrichment of NF-κB regulatory elements in HybHep-accessible chromatin without corresponding transcriptional activation supports a model of epigenetic priming, in which the regulatory landscape is pre-configured for rapid signal-driven transcriptional responses upon injury. Our GSEA results are consistent with the predominant enrichment of NF-κB motifs among injury-gained DARs identified in Figure 2. Altogether, HybHeps encode a permissive chromatin landscape in homeostasis, characterized by active enhancer configurations at injury-responsive loci and selective epigenetic priming for NF-κB–mediated inflammatory activation. These findings raise the question of whether the chromatin mechanisms that enable injury-responsive transcription are distinct from those that stabilize HybHep identity.

### Injury-responsive transcription is driven primarily by accessibility remodeling, whereas state-associated and shared programs are stabilized by histone modifications

Having established that HybHeps encode a permissive enhancer landscape at injury-responsive loci relative to cHeps (**Figure 3**), we next asked how distinct transcriptional programs within HybHeps are encoded at the chromatin level during injury and in the cell state context. To determine whether chronic injury and hepatocyte cell state activate distinct or overlapping transcriptional programs, we compared statistically significant log2 fold changes of differentially expressed genes across conditions. Plotting injury-associated versus state-associated expression changes revealed a structured quadrant distribution defining three principal gene classes: injury-responsive, state-associated, and shared (**Figure 4A, Supplemental Figure 4A**).

**Figure 4.**
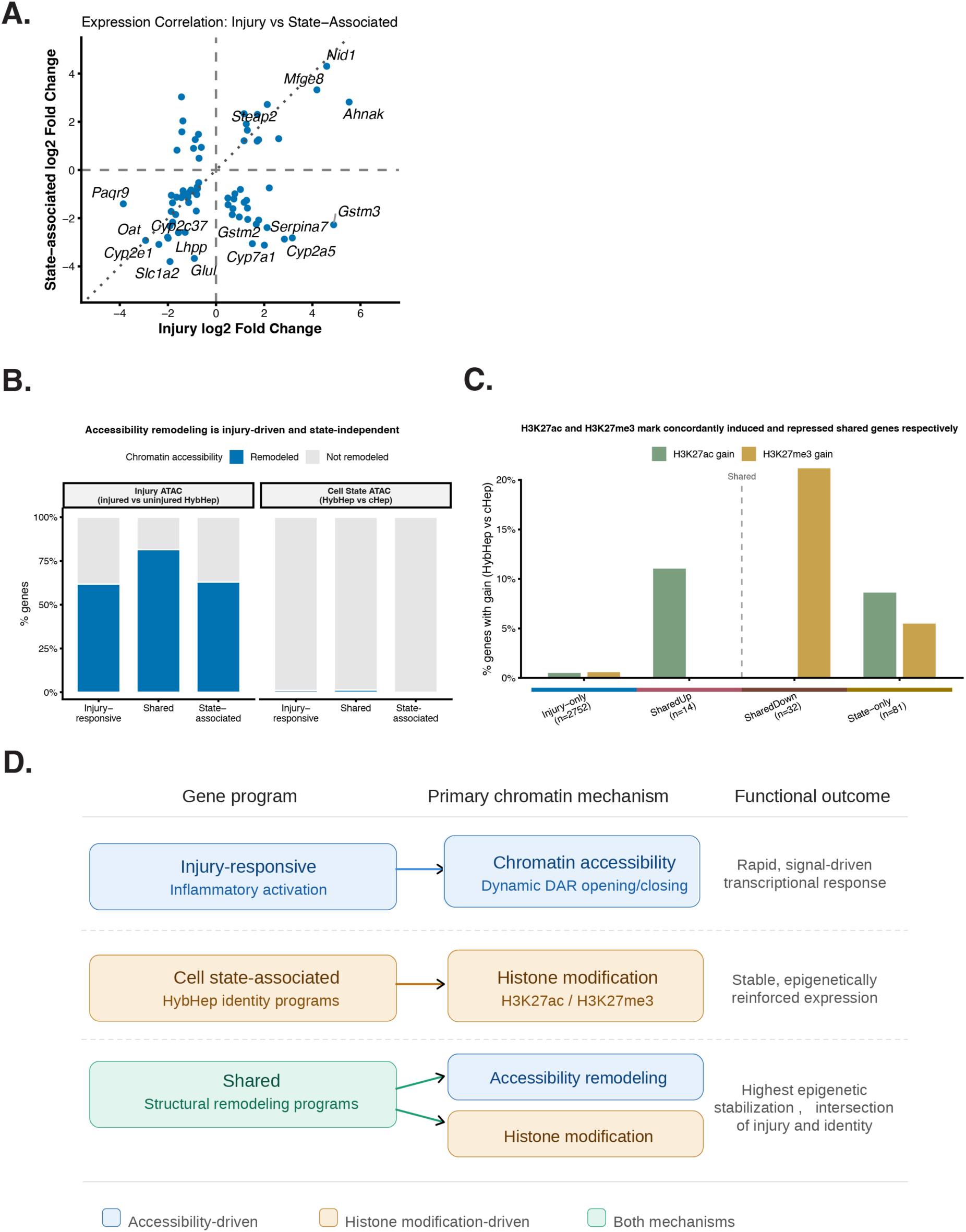
Chromatin regulatory logic underlies injury-responsive and state-stabilized transcriptional programs. **(A)** Scatter plot of log2 fold changes for genes differentially expressed in both injury and state contexts. Quadrants denote concordant or discordant regulation, highlighting genes with shared, injury-specific, or state-biased transcriptional responses. **(B)** Fraction of genes exhibiting chromatin accessibility changes (open or closed) in response to injury or cell state, stratified by differential expression program (injury-only, shared, state-only). Injury-responsive genes show a markedly higher frequency of ATAC remodeling compared with state-only genes, whereas state-associated accessibility changes are rare across all programs. Bars are grouped by transcriptional program and colored accordingly. **(C)** Bar plot showing the percentage of genes in each program class with significant gains in H3K27ac or H3K27me3 in HybHeps relative to cHeps (DESeq2, adjusted P < 0.05, log2FC > 0.5; peaks assigned to nearest gene). The shared program is split into concordantly induced (SharedUp, n=14) and concordantly repressed (SharedDown, n=32) subsets. H3K27ac gain is preferentially associated with SharedUp genes, while H3K27me3 gain is concentrated among SharedDown and state-associated genes, suggesting mark-appropriate histone remodeling rather than bivalency at shared loci. Injury-responsive genes show minimal histone remodeling in either direction. **(D)** Schematic summarizing distinct chromatin regulatory logic, in which injury-responsive transcription is encoded primarily through dynamic chromatin accessibility, while state-associated and shared programs are stabilized through histone remodeling.

Shared genes exhibited concordant regulation in both contexts and were enriched for regulators of cytoskeletal organization, extracellular matrix interaction, and membrane remodeling, including *Nid1*, *Spp1*, *Mfge8*, *Anxa5*, and *Ahnak*, consistent with a common structural remodeling program. Concordantly repressed genes were dominated by canonical hepatocyte metabolic pathways, including cytochrome P450 family members (*Cyp2e1*, *Cyp2c38*, *Cyp2c37*), bile acid metabolism genes (*Cyp27a1*, *Hsd3b7*), and glutamine metabolism genes (*Glul*, *Oat*), reflecting suppression of mature hepatocyte function across both states. In contrast, injury-responsive genes were preferentially induced during injury and included stress-response and detoxification programs (*Hamp*, *Aldh1a1*, *Gstm1/2/3*, *Ephx1*). State-associated genes were selectively enriched in HybHeps and included metabolic enzymes (*Hal*, *Sds*, *Uroc1*, *Fbp1*), regulators of cytoskeletal dynamics (*Arhgap6*, *Arhgap10*), and growth factor signaling components (*Inhba*), suggesting engagement of programs associated with stabilization of a specific cell state rather than acute inflammatory signaling.

We next asked whether these transcriptional program classes are associated with distinct chromatin accessibility changes. To address this, we intersected each gene class with two independent ATAC-seq differential accessibility analyses: an injury comparison (MetHeps versus HybHeps, n=4 per group) and a cell state comparison (HybHep versus cHep, n=3 per group). A gene was classified as accessibility-remodeled if its nearest DAR reached significance in the respective comparison (adjusted P < 0.1, |log2FC| > 0.5). In the injury ATAC comparison, a substantial fraction of genes across all three program classes showed chromatin accessibility changes, reflecting the broad scope of injury-induced remodeling (**Figure 4B, left**). By contrast, virtually no genes in any program class showed significant accessibility changes in the cell state comparison (**Figure 4B, right**), indicating that the transcriptional differences between HybHeps and cHeps — including both state-associated and shared programs — are maintained largely without widespread differences in steady-state chromatin accessibility. Together these findings suggest that injury drives acute accessibility remodeling across multiple gene programs, whereas state-associated transcriptional identity is encoded through mechanisms other than differential chromatin accessibility.

Despite the lack of widespread accessibility changes at state-associated loci, histone modification patterns revealed a distinct regulatory logic. To assign histone modification status to genes, significant H3K27ac and H3K27me3 peaks from the HybHep versus cHep CUT&Tag comparison were mapped to their nearest gene using ChIPseeker (adjusted P < 0.05, log2FC > 0.5). Gains in H3K27ac and H3K27me3 in HybHeps relative to cHeps were selectively enriched among state-associated and shared gene sets, while injury-responsive genes showed minimal histone remodeling (**Figure 4C**). Importantly, splitting the shared gene set by expression direction revealed that H3K27ac gain was preferentially associated with concordantly induced shared genes (SharedUp, n=14), while H3K27me3 gain was concentrated among concordantly repressed shared genes (SharedDown, n=32). This pattern indicates that the apparent co-enrichment of both marks at shared genes reflects the mixed transcriptional directionality of this class rather than bivalency at individual loci– H3K27ac consolidates, activated programs while H3K27me3 stabilizes repressed programs, independent of widespread chromatin accessibility remodeling. We propose a model in which injury-responsive transcription is mediated primarily through dynamic chromatin accessibility changes, whereas state-associated and shared transcriptional programs are reinforced through selective histone modification within a context of open chromatin, consistent with stabilization of cell-state identity (**Figure 4D**).

### Human liver atlas analysis reveals a SOX9-associated injury-responsive hepatocyte state

To determine whether the *Sox9*-associated injury-responsive hepatocyte state identified in mice is represented in the human liver, we analyzed the GepLiver human liver single-cell atlas^11^, which encompasses hepatocytes from both healthy and diseased livers, restricted to adult non-malignant hepatocytes (Normal n=65, MASLD n=265, Viral Hepatitis n=28, Cirrhosis n=20). Dimensional reduction and UMAP embedding of this hepatocyte subset revealed a structured manifold reflecting transcriptional heterogeneity across disease contexts. SOX9 expression was confined to a rare subset of hepatocytes distributed across the manifold (**Figure 5A**). The low frequency of SOX9-expressing hepatocytes is consistent with both the known rarity of periportal SOX9⁺ hepatocytes *in vivo* and the inherent dropout characteristics of scRNA-seq for lowly expressed genes, which may result in underrepresentation of cells with biologically meaningful but low-level SOX9 expression.

**Figure 5.**
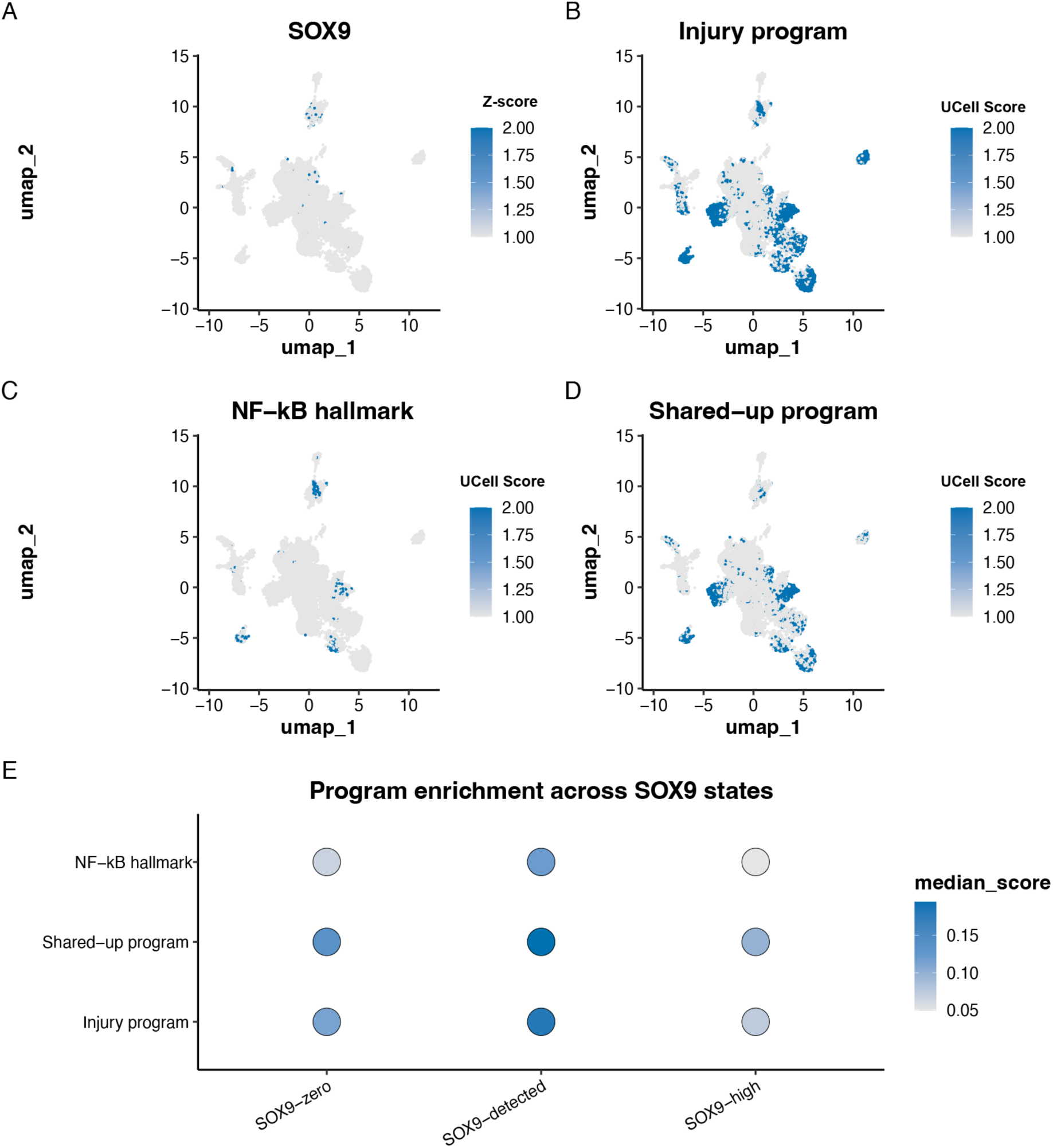
Human liver atlas analysis identifies a SOX9-associated injury-responsive hepatocyte state. **(A–D)** UMAP embeddings of adult non-malignant hepatocytes from the GepLiver human liver single-cell atlas, colored by normalized SOX9 expression (A) and UCell scores for the injury-responsive program (B), NF-κB hallmark gene set (C), and shared-upregulated program (D). Gray cells have zero or below-threshold expression or score. Dimensional reduction was performed on the hepatocyte subset following Harmony batch correction. **(E)** Dot plot showing median UCell scores for the injury-responsive, shared-upregulated, and NF-κB hallmark programs across hepatocytes stratified by SOX9 expression state (SOX9-zero, SOX9-detected, SOX9-high). Dot size encodes median score magnitude. Hepatocytes with detectable SOX9 expression show the highest program enrichment, while SOX9-high cells show modestly attenuated scores relative to SOX9-detected cells, suggesting heterogeneity within the SOX9-positive compartment.

Projection of mouse-derived injury-responsive, shared-upregulated, and NF-κB hallmark gene programs onto the human hepatocyte manifold revealed enrichment that was regionally distributed rather than uniform, with high-scoring cells partially overlapping with areas of detectable SOX9 expression (**Figure 5B–D**). This spatial correspondence is consistent with a conserved association between SOX9 expression and inflammatory program activity in human hepatocytes, though the broader distribution of program enrichment indicates that injury-associated transcriptional states are not exclusively restricted to SOX9-expressing cells.

To more directly assess the relationship between SOX9 expression level and program enrichment, hepatocytes were stratified into three expression states based on normalized SOX9 expression: SOX9-zero (no detectable expression, n = 14,972), SOX9-detected (expression above zero but below the top 10th percentile threshold, n = 449), and SOX9-high (normalized expression > 2.99, n = 50) (**Supplemental Figure 5A**). Median UCell scores were then compared across these groups (**Figure 5E**). SOX9-detected cells showed the highest median enrichment for all three programs, injury-responsive, NF-κB hallmark, and shared-upregulated, exceeding both SOX9-zero and SOX9-high cells. The observation that SOX9-high cells showed modestly lower program enrichment than SOX9-detected cells suggests that transcriptional program activity is not simply proportional to SOX9 expression dosage and may instead reflect functional heterogeneity within the SOX9-expressing hepatocyte compartment, or a non-linear relationship between SOX9 levels and downstream inflammatory activation. Notably, the relative rarity and attenuated inflammatory program enrichment of SOX9-high cells raises the possibility that this population may be phenotypically closer to biliary epithelial cells and further along a hepatocyte-to-biliary transdifferentiation trajectory, consistent with the known role of SOX9 in biliary specification (^2,3^). Together, these findings support partial conservation of the mouse-defined SOX9-associated injury-responsive hepatocyte state in the human liver, suggesting that inflammatory program enrichment in human hepatocytes is most pronounced at intermediate rather than maximal SOX9 expression levels.

## Discussion

### A chromatin-based dichotomy parses injury-responsive and state-stabilized hepatocyte states

Hepatocytes exhibit remarkable plasticity in response to injury, yet the regulatory logic that distinguishes a chronic injury response from stable cell state has remained unclear. Through integrative multi-omic profiling of cHeps, HybHeps, and MetHeps, we define a chromatin-based dichotomy in which injury-responsive transcription and state-associated gene regulation are mediated by fundamentally distinct epigenetic mechanisms.

Injury-responsive genes were characterized by widespread chromatin accessibility remodeling and enrichment for NF-κB, AP-1, and STAT motifs, consistent with rapid, signal-dependent activation of pre-existing regulatory elements within *Sox9+* hepatocytes. This pattern is consistent with a higher barrier to pro-inflammatory transcription in MetHeps relative to HybHeps, where accessibility differences are most pronounced and may reflect a permissive chromatin state that facilitates inflammatory transcription factor binding^10,12^. We find that HybHeps and cHeps exhibit broadly similar accessibility landscapes, including differential H3K27ac and H3K27me3, representing the predominant epigenetic distinction between these states. H3K27ac marks active enhancers and is associated with transcriptional activation^13^, while H3K27me3 enforces facultative repression and is central to Polycomb-mediated stabilization of state identity^14^. This suggests that the transition between hybrid and conventional hepatocyte identity does not require large-scale chromatin remodeling, but may instead be fine-tuned by histone post translational modifications that reinforce subtle regulatory differences across shared open chromatin regions, permitting transcription factor engagement without necessitating accessibility turnover.

Importantly, genes shared between injury and cell state programs displayed the strongest enrichment for histone remodeling and, notably, did not show corresponding differences in chromatin accessibility, consistent with their regulation occurring within a constitutively open chromatin landscape that is further refined by histone modification state rather than accessibility gating. The opposing behavior of SharedUp and SharedDown modules — in which injury-associated activation is mirrored by coordinated repression of metabolic identity genes — argues against nonspecific transcriptional activation and reinforces the structured organization of injury and cell state programs that distinguish HybHeps and cHeps. Formal characterization of the SharedDown module across human disease contexts represents a priority for subsequent analysis.

Projection of these mouse-defined programs onto a human single-cell hepatocyte atlas¹¹ is consistent with partial conservation of this chromatin architecture. SOX9-expressing hepatocytes in human liver showed preferential engagement of the injury-associated inflammatory module and concordant induction of shared genes, suggesting reduced activity of concordantly repressed metabolic identity genes in this population. Program enrichment was highest among SOX9-detected cells (those with low but detectable SOX9 expression) rather than among SOX9-high cells, suggesting that inflammatory program activity is not simply proportional to SOX9 expression dosage and may reflect a non-linear relationship between SOX9 levels and downstream transcriptional activation. Although the magnitude of program enrichment varies across disease contexts, the directional consistency across inflammatory, shared activation, and identity repression modules is consistent with partial conservation of the injury–state dichotomy across species.

These findings refine current models of hepatocyte plasticity. Rather than representing a binary conversion between hepatocyte and biliary identities^5,7^, HybHeps appear to occupy a regulated transitional state defined by accessibility-driven inflammatory activation and selective epigenetic stabilization of remodeling programs. In this framework, injury-responsive transcription is primarily governed by rapid chromatin accessibility changes^9^, whereas state-associated stabilization is reinforced through histone modification–mediated chromatin consolidation. SOX9-associated regulatory pathways may participate in this stabilization process, but the predominant instructive drivers of injury remodeling appear to be inflammatory transcription factors.

Conceptually, these data argue that hepatocyte identity and injury response is encoded by separable and hierarchically organized chromatin layers that differentially govern inflammatory activation and state stabilization. Accessibility remodeling permits rapid inflammatory activation, while histone modifications consolidate a subset of shared and state programs. The balance between these layers may determine whether hepatocytes revert to homeostasis or progress toward sustained phenotypic alteration in chronic disease. More broadly, this injury–state dichotomy may represent a generalizable principle of epithelial plasticity^12^, in which stress responses and stabilized identity shifts are encoded by separable chromatin mechanisms. Future studies leveraging temporal perturbation and state tracing will be required to determine how these regulatory modules are dynamically coordinated during disease progression and regeneration.

## Methods

### RNA-SEQ PROCESSING

The Nextflow^15^ workflow manager was implemented to build and maintain a bespoke RNA-SEQ preprocessing pipeline that we’ve previously published^16^. In this pipeline, raw reads were inspected for adapter and overrepresented sequences, as well as GC content per sample with FastQC (v1.11). FastQC results per sample were aggregated for visualization with the MultiQC^17^. The samples that passed quality control are aligned to the GRCm38.p6 genome using STAR (v2.7.7a)^18^ and transcripts were quantified using Salmon^19^ (1.8.0). SAMtools^20^ was used to convert raw reads to SAM and BAM formats for post-alignment processing. Picard (v2.23.4) CollectMultipleMetrics was run on Bam files to collect classes of quality control metrics describing base distribution. Picard CollectRnaSeqMetrics was run on BAM files to produce RNA alignment metrics. Picard MarkDuplicates was run on BAM files to tag duplicate reads. Tximeta (v1.44.0)^21^ was used to assign transcriptome metadata to the counts matrix.

### DIFFERENTIAL GENE EXPRESSION MODEL: Injured v Uninjured

The DESeq2^22^ negative binomial generalized linear model was used for differential gene expression analysis between DDC-treatment groups. We observed a sample preparation batch effect by PCA in the dataset so we included batch as a covariant in the model: design = ∼ batch + Treatment. Model prediction output statistics resulted from a comparison of injured to uninjured samples (n = 4 per group).

### DIFFERENTIAL GENE EXPRESSION MODEL: EGFP+ v EFGP−

The DESeq2^22^ negative binomial generalized linear model was used for differential gene expression analysis between EGFP-treatment groups. We observed a mouse effect by PCA but did not include this as a model covariate because it minimally impacted treatment results (data not shown). We also observed a sample preparation batch effect in PC1 and included this as a continuous covariate in the model: design = ∼ PC1 + Treatment. Model prediction output statistics resulted from a comparison of EGFP+ to EGFP− samples (n = 3 per group). We identified 42 genes significantly upregulated (log2 (FC) > 0), 115 genes significantly downregulated (log2 (FC) < 0) and 9,147 low counts (adjusted p value< 0.1).

### ATAC-SEQ DATA PROCESSING

The Nextflow workflow manager was implemented to build and maintain a bespoke ATAC-seq preprocessing pipeline. Raw reads were trimmed and quality-checked using Trim Galore (v0.6.7) with the --fastqc flag for paired-end input. Trimmed reads were aligned to the mouse genome (mm10) using Bowtie2^23^ with default settings (v2.5.4). Resulting SAM files were processed and duplicate-marked with Picard (v2.20.0) MarkDuplicates, and insert size metrics were collected using Picard CollectInsertSizeMetrics. Peaks were called using MACS (v2.2.9.1)^24^. FRiP scores per sample were calculated using deepTools (v3.5.4)^25^ plotEnrichment -outRawCounts and ranged from 0.13–0.30. MSPC (v6.1.0) ^26^ was run using the command line interface with the parameters: -r bio -w 1e-6 -s 1e-14 -c 3 -m lowest -d 4 for joint analysis of weak peaks across replicates. Reads in peaks were counted using csaw (Bioconductor v1.38.0)^27^ and the resulting count matrix was used for downstream analysis. Peak sets for each condition were annotated using ChIPseeker (v1.40.0)^28^ annotatePeak function and the UCSC Genome Browser knownGene (mm10) table. Peaks mapped to the mitochondrial genome and overlapping with the mm10 ENCODE blacklist^29^ were excluded.

### DIFFERENTIAL REGION ACCESSIBILITY MODEL: Injured v Uninjured

DESeq2^24^ was used for differential region accessibility analysis of 57,349 peaks. We observed a sample batch preparation effect and included this as a covariate in the DESeq2 model: design = ∼Batch + Treatment. Model prediction output statistics resulted from a comparison of injured versus uninjured hybrid hepatocytes (n = 4 per group). We identified 13,846 (24%) regions significantly upregulated (log2 (FC) > 0), 13,273 (23%) regions significantly downregulated (log2 (FC) < 0) and 4 count outliers (0.007%).

### ATAC DIFFERENTIAL REGION ACCESSIBILITY MODEL: EGFP+ v EFGP−

Of the 48,065 peaks called across samples, peaks (read count <10) that were not present in 2/3 samples per condition were inspected and excluded from differential region accessibility modeling. Peaks mapped to the mitochondrial genome (2 peaks) and overlapped with the mm10 ATAC blacklist (112 peaks) were also excluded. DESeq2 was used for differential region accessibility analysis of the remaining peaks. We observed a mouse effect by PCA but did not include this as a model covariate because it minimally impacted treatment results (plots not shown). We also observed a sample batch preparation effect by PCA in PC1 and included PC1 as a covariate in the model: design = ∼PC1 + condition. Model prediction output statistics resulted from a comparison of SOX9+/EGFP+ to SOX9-/EGFP− samples (n = 3 per group).

### CUT&Tag PROCESSING

Reads were aligned to the *mm10* reference genome using Bowtie2. Average fragment size per assay was determined using Bedtools. For H3K27ac, peaks were called using MACS2 (v2.2.9.1) with the following parameters: -f BAMPE -q 0.05 --keep-dup auto --nomodel --extsize 304 -g mm. For H3K27me3, broad peaks were called with parameters: -f BAMPE -g mm --broad. Consensus peaks within groups were identified using MSPC with Fisher’s method and the following settings: -r bio -w 1e-4 -s 1e-5 -c 2. Peaks mapping to the mitochondrial genome or overlapping the *mm10* CUT&RUN blacklist were excluded.

For H3K27me3, read fragments within peaks were quantified using csaw::regionCounts on the aligned BAM files. To estimate a global background signal, low-abundance windows were filtered out using csaw::windowCounts, and the resulting normalization factors were used in place of DESeq2 size factors. FRiP scores were calculated using deepTools (v3.5.4) with plotEnrichment --outRawCounts, and ranged from 9 to 12%.

### CUT&TAG DIFFERENTIAL REGION ACCESSIBILITY MODELS: EGFP+ v EGFP-

Peaks that were not present in 2/3 samples per condition were inspected and excluded from differential region accessibility modeling. DESeq2 was used for differential region accessibility analysis using the model: design =∼condition. Model prediction output statistics resulted from a comparison of SOX9+/EGFP+ to SOX9-/EGFP− samples (n = 3 per group).

### GENE SET ENRICHMENT ANALYSIS: RNA-SEQ

The package fgsea^30^ (v1.30.0) was used to run GSEA. Gene lists ranked by DESeq2 results statistics were used as gene set inputs. GSEA was run with default settings, without collapsing, and with the gene set as the permutation type. The leading-edge analysis function was used to determine the significance of gene set enrichment.

### GENE SET ENRICHMENT ANALYSIS: ATAC-SEQ and CUT&Tag

A consensus peak set was derived from uninjured and injured or EGFP+ and EGFP− peaks using MSPC. For each peak, the log2FC, p-value, and adjusted p-value was calculated. Annotated peaks were ranked based on the log2FC prior to Gene Set Enrichment Analysis. The ranked list of gene symbols is used as input to perform Gene Set Enrichment Analysis with the fgseaMultilevel function of the R package fgsea(v1.30.0)^31^.

### DATA VISUALIZATION

All box plots, volcano plots, and scatter plots were generated in R using ggplot2 (v4.0.1)^32^. Volcano plots display ApeGLM-shrunken^33^ log2 fold changes on the x-axis and −log10 adjusted p-values on the y-axis, with significance thresholds indicated by dashed lines. Gene labels were added using ggrepel (v0.9.6) to minimize label overlap. Box plots display variance-stabilized (VST) normalized counts per sample, with individual data points overlaid. Scatter plots of ATAC-seq versus RNA-seq fold changes display the maximum ATAC signal per gene. Genome browser tracks were generated using plotgardener and display mean-normalized signal across biological replicates (n=3-4).

### DIGITAL GENOMIC FOOTPRINTING ANALYSIS

Tn5 transposase cut site signal is hindered by protein-DNA interactions which results in decreased signal strength in high-signal genomic regions (genomic footprints)^29^. The TOBIAS^34^ digital genomic footprinting framework reasons that correcting for Tn5 bias uncovers 50%+ of measurable transcription factor-DNA binding. The TOBIAS^35^ Snakemake workflow was applied to predict transcription factor binding across ATAC-Seq sample conditions. The workflow uses bam-formatted ATAC-seq reads, transcription factor mononucleotide position weight matrices (PWMs) and sequence annotation as inputs. PWMs were queried using pyJASPAR and then written to JASPAR format and saved as .jaspar. During processing, replicate BAMs are joined per condition to gain sequencing depth required for peak annotation, Tn5 bias correction, motif binding position estimation, and statistical comparison of motif occurrences.

### HUMAN LIVER SINGLE-CELL ATLAS

To determine whether the SOX9-associated injury-responsive hepatocyte state identified in mouse was represented in human liver, we analyzed a published human liver single-cell RNA-seq atlas (GepLiver)^11^. A preprocessed Seurat object was loaded into R and restricted to cells annotated as hepatocytes (Cell_Type == “Hepatocyte”). To focus on adult nonmalignant hepatocytes, cells corresponding to tumor-associated or fetal-like states were excluded based on reconstructed phenotype labels derived from orig.ident, and the analysis was further restricted to cells annotated as Hepa_Normal in the Cell_Subtype metadata field.

SOX9 expression was extracted from the normalized RNA assay (layer = “data”), and hepatocytes were stratified into three expression states: SOX9-zero, SOX9-detected, and SOX9-high. Cells with zero normalized SOX9 expression were classified as SOX9-zero. Among cells with detectable SOX9 expression, the top 10% of SOX9-expressing cells were classified as SOX9-high, while the remaining SOX9-positive cells were classified as SOX9-detected.

To visualize the hepatocyte state manifold, dimensional reduction was recomputed on the filtered subset using principal component analysis (PCA) followed by Harmony correction for dataset-level effects (dataset_short) and UMAP embedding. Mouse-derived transcriptional programs were projected onto the human hepatocyte atlas using UCell. Program scores were calculated for the injury-responsive, shared-upregulated, and NF-κB hallmark gene sets after restricting each signature to genes detected in the human dataset. UMAPs were colored by normalized SOX9 expression or UCell score, and program enrichment across SOX9 states was summarized using median UCell scores.

All analyses were performed in R using Seurat^36^, Harmony^37^, and UCell^38^.

## Author contributions

JAB, FH, JR, and ADG designed experiments and analyzed data.

JAB, FH, and ADG conducted experiments.

JAB wrote the manuscript.

JAB, JR, and ADG edited the manuscript.

---

**Supplemental Figure 1.**
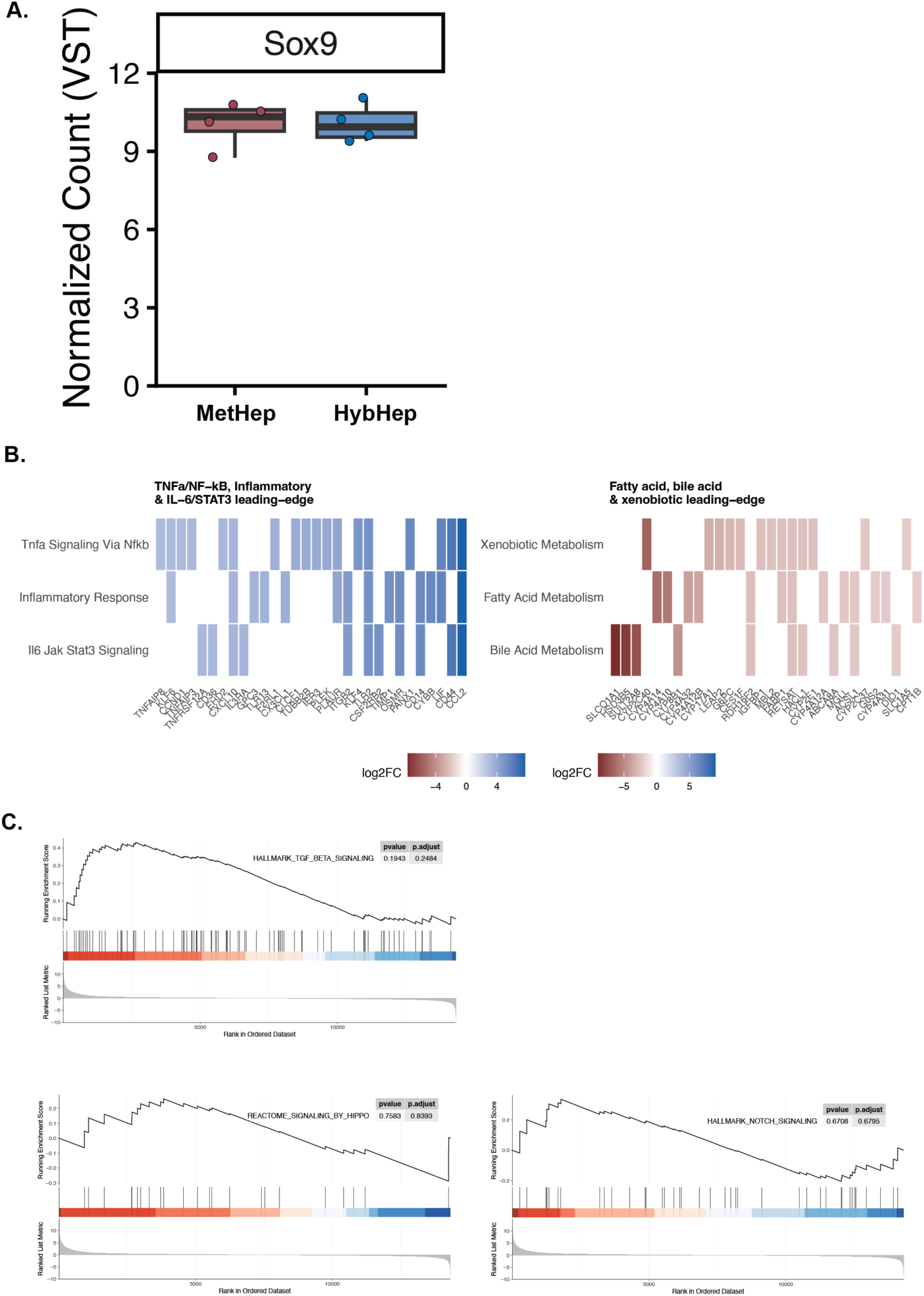
Transcriptional features of injury-induced hybrid hepatocytes. **(A)**Variance-stabilized RNA-seq counts for *Sox9* in control and DDC-treated hybrid hepatocytes. **(B)** GSEA of Hallmark gene sets highlighting signaling pathways commonly associated with hepatoductal transdifferentiation. **(C)** Leading-edge gene statistics for the most significantly enriched gene sets.

**Supplemental Figure 2.**
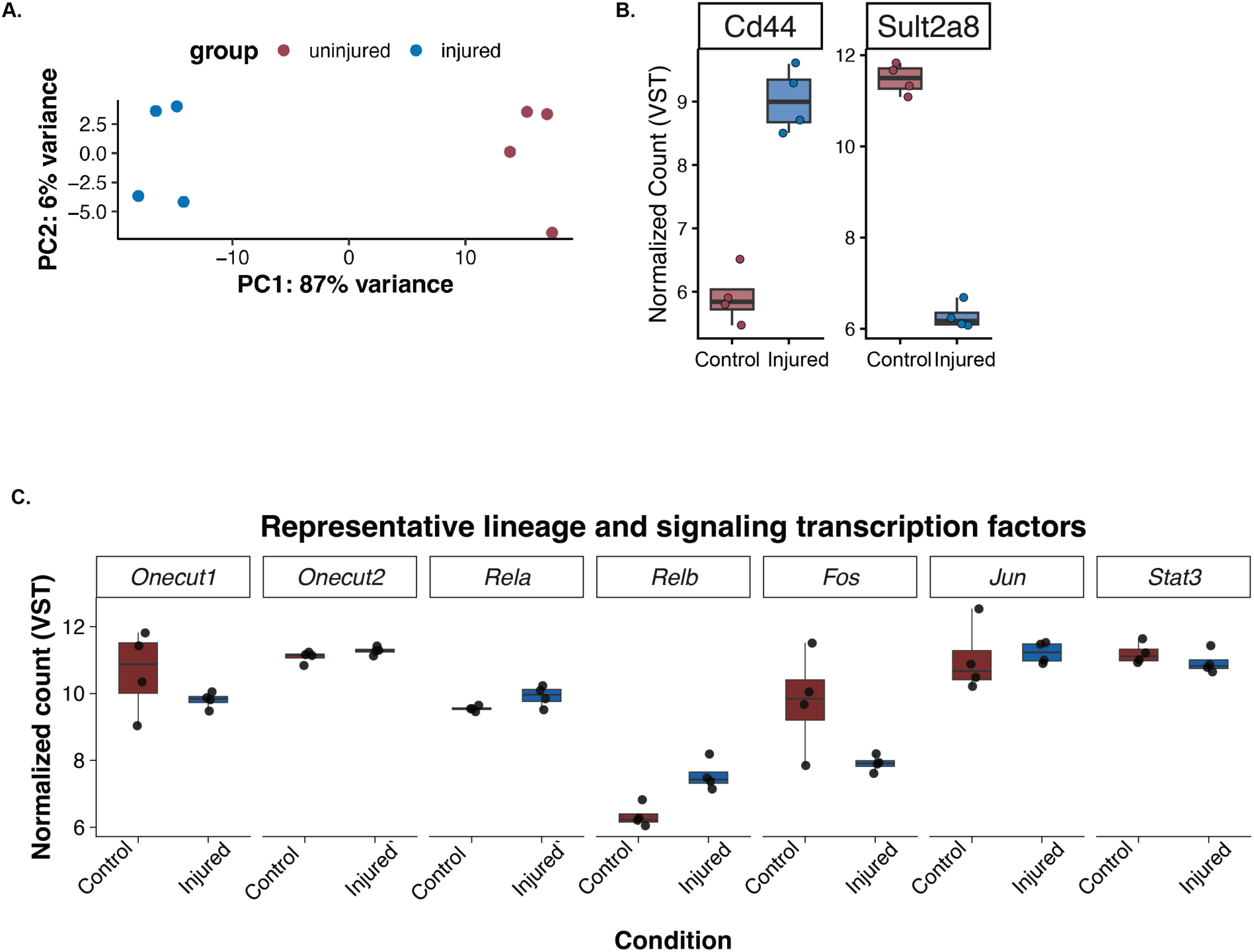
Global chromatin accessibility changes in metaplastic hepatocytes. **(A)** Principal component analysis of ATAC-seq samples from control and DDC-injured hybrid hepatocytes (n=4 per group). PC1 accounts for 87% of total variance and separates samples by injury status, suggesting robust and reproducible injury-associated chromatin remodeling. **(B)** VST-normalized expression of representative inflammatory signaling and hepatocyte state genes, *Cd44* and *Sult2a8* (n = 4). **(C)** VST-normalized expression of representative inflammatory signaling and hepatocyte state transcription factors in uninjured and DDC-treated hybrid hepatocytes (n=4 per group). *Onecut1* is significantly downregulated following injury, consistent with predicted loss of ONECUT occupancy at injury-repressed chromatin regions. *Onecut2* shows a modest increase in the injured condition; the significance of this divergent response relative to *Onecut1* is unclear and may reflect distinct transcriptional roles of the two paralogs.

**Supplemental Figure 3.**
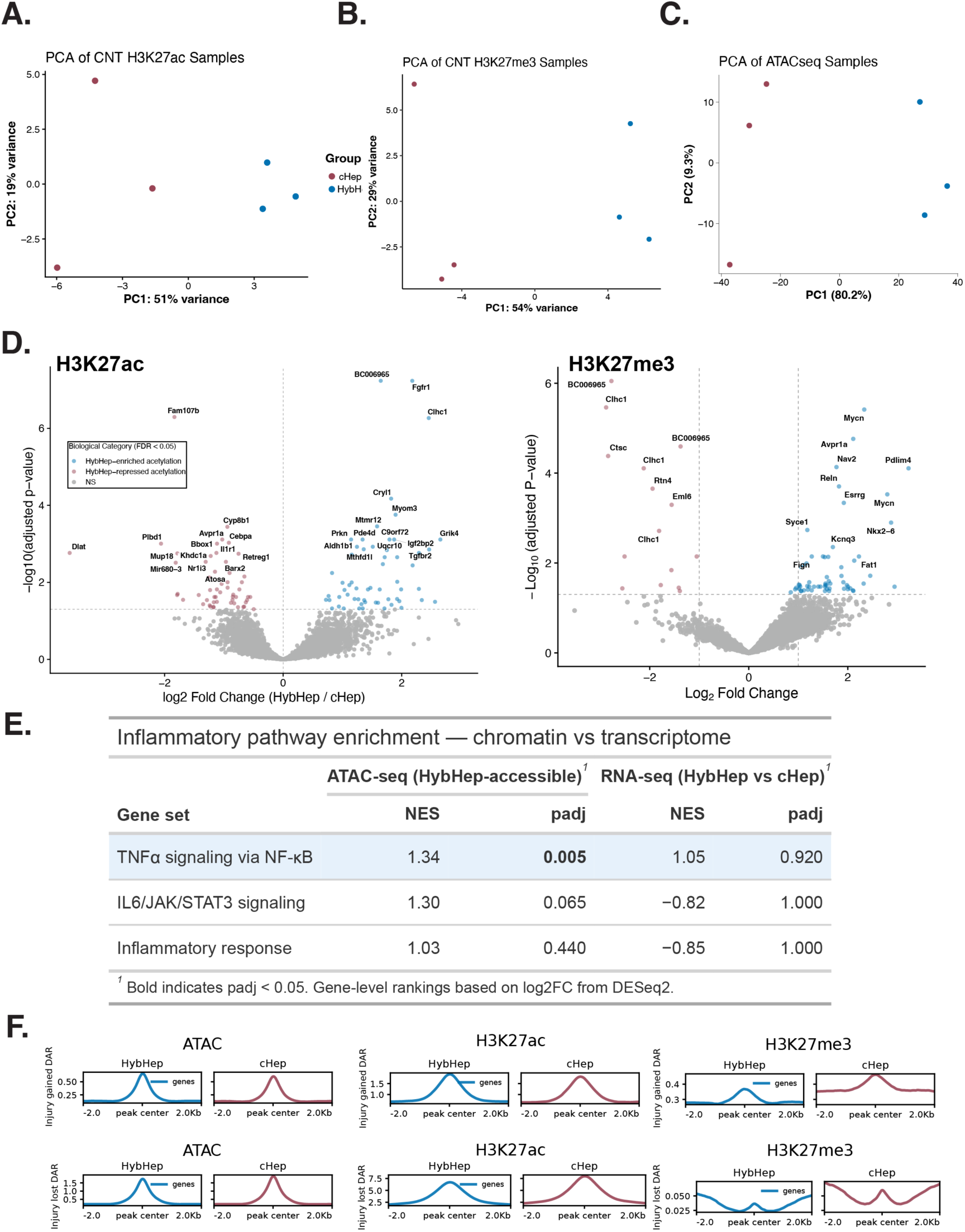
Quality control and differential epigenomic profiling of hybrid and conventional hepatocytes. **(A)** Principal component analysis of CUT&Tag H3K27ac samples from cHep and HybHep populations (n=3 livers). PC1 accounts for 51% of total variance and separates samples by cell type. **(B)** Principal component analysis of CUT&Tag H3K27me3 samples (n=3 livers). PC1 accounts for 54% of total variance and separates samples by cell type. **(C)** Principal component analysis of ATAC-seq samples from cHep and HybHep populations (n=3 livers). PC1 accounts for 87% of total variance and separates samples by cell type. **(D)** Volcano plots showing log₂ fold change and adjusted p-values for differential H3K27ac (left) and H3K27me3 (right) enrichment between HybHeps and cHeps. HybHep-enriched and HybHep-repressed regions are highlighted (adjusted P < 0.05). **(E)** Gene set enrichment analysis of genes linked to HybHep-enriched accessible chromatin regions against MSigDB Hallmark gene sets. Bars show normalized enrichment scores (NES) for the three most relevant inflammatory pathway gene sets. TNFα signaling via NF-κB is significantly enriched (NES = 1.34, padj = 0.005), while IL6/JAK/STAT3 signaling shows a positive trend (NES = 1.30, padj = 0.065) and the hallmark inflammatory response gene set is not significantly enriched (NES = 1.03, padj = 0.44). Gene-level ranking was based on ATAC-seq log2 fold change (HybHep/cHep). Tracks display mean signal across biological replicates (n=3). Signal ranges are matched within each assay and indicated in brackets. **(F)** Meta-profile plots showing mean signal ±2kb showing ATAC-seq, H3K27ac, and H3K27me3 signal centered ±2 kb from the midpoint of injury-gained (top) and injury-lost (bottom) accessible regions, shown for HybHep and cHep samples. At injury-gained DARs, HybHeps show higher ATAC-seq and H3K27ac signal relative to cHeps, with comparatively lower H3K27me3, consistent with a permissive baseline enhancer configuration at injury-responsive loci. At injury-lost DARs, HybHeps show reduced H3K27ac relative to cHeps, while H3K27me3 levels are similarly low in both cell types, indicating that transcriptional repression at hepatocyte metabolic gene loci is not associated with Polycomb-mediated H3K27me3 deposition. Signals represent mean-normalized values across biological replicates (n=3 livers per group). Rows are sorted by HybHep ATAC-seq signal at injury-associated DARs and carried through to all subsequent panels.

**Supplemental Figure 4A.**
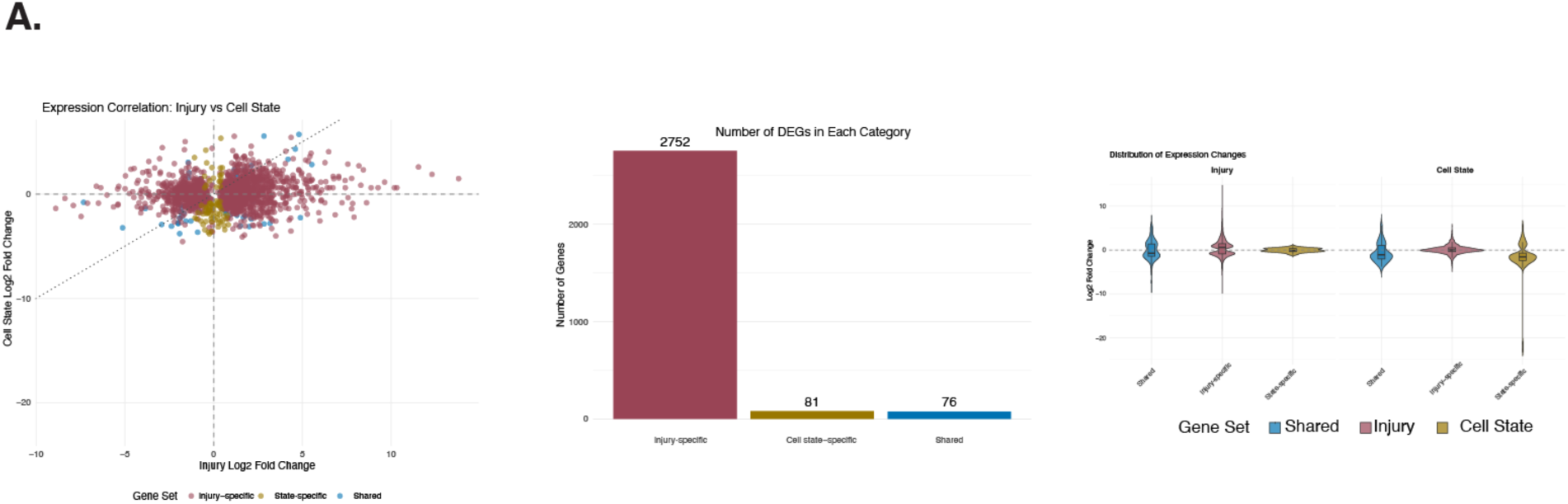
Gene program classification and chromatin priming of shared transcriptional programs. Left: Scatter plot of injury versus state log2 fold changes for all differentially expressed genes, colored by program class (injury-only, state-only, shared). A total of 2,752 injury-only, 81 state-only, and 76 shared differentially expressed genes were identified. Center: Bar chart showing the number of genes in each program class. Right: Violin plots showing the distribution of log2 fold changes for each gene class in the injury (left) and state (right) comparisons, confirming directional consistency within each class.

**Supplemental Figure 5A.**
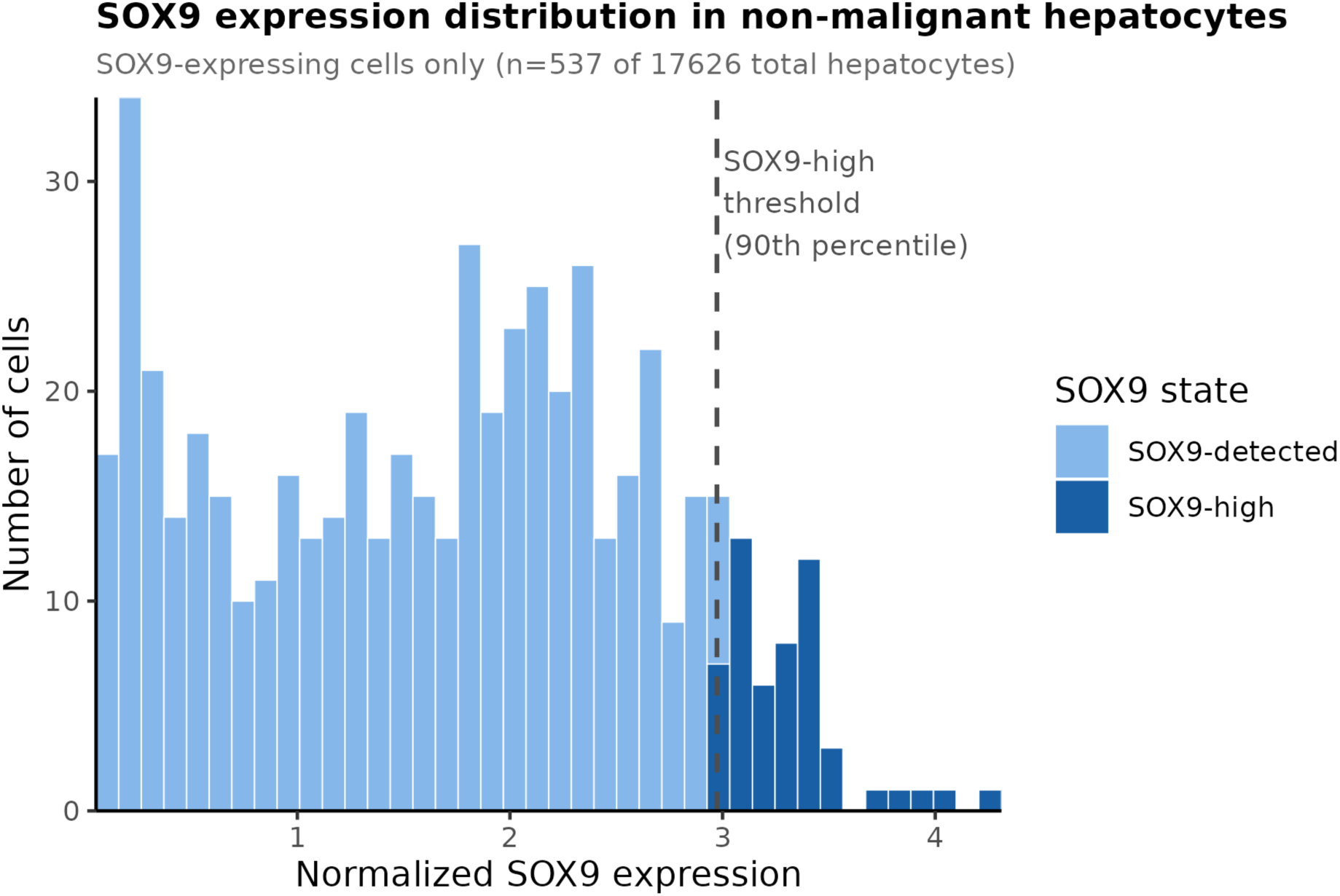
SOX9 expression distribution in adult non-malignant hepatocytes from the GepLiver human liver atlas. Histogram of normalized SOX9 expression among SOX9-expressing hepatocytes (n = 499 of 15,471 total cells). Cells are colored by SOX9 expression state: SOX9-detected (light blue, expression > 0 but below threshold) and SOX9-high (dark blue, top 10th percentile of expressing cells). The dashed line indicates the SOX9-high cutoff (normalized expression = 2.99). SOX9-zero cells (n = 14,972) are excluded from this display as they have no detectable SOX9 expression.

